# Monoubiquitination by the Fanconi Anemia core complex locks FANCI:FANCD2 on DNA in filamentous arrays

**DOI:** 10.1101/862805

**Authors:** Winnie Tan, Sylvie van Twest, Andrew Leis, Rohan Bythell-Douglas, Vincent J. Murphy, Michael Sharp, Michael W Parker, Wayne M Crismani, Andrew J. Deans

## Abstract

FANCI:FANCD2 monoubiquitination is a critical event for replication fork stabilization by the Fanconi anemia (FA) DNA repair pathway. It has been proposed that at stalled replication forks, monoubiquitinated-FANCD2 serves to recruit DNA repair proteins that contain ubiquitin-binding motifs. Here we have reconstituted the FA pathway *in vitro* to study functional consequences of FANCI:FANCD2 monoubiquitination. We report that monoubiquitination does not promote any specific exogenous protein:protein interactions, but instead stabilizes FANCI:FANCD2 heterodimers on dsDNA. This locking of FANCI:FANCD2 complex on DNA requires monoubiquitination of only the FANCD2 subunit. We further show that purified monoubiquitinated FANCI:FANCD2 forms filament-like arrays on long dsDNA using electron microscopy. Our results reveal how monoubiquitinated FANCI:FANCD2 is activated upon DNA binding and present new insights to potentially modulate monoubiquitinated FANCI:FANCD2/DNA filaments in FA cells.

## Introduction

Fanconi anemia (FA) is a devastating childhood syndrome that results in bone marrow failure, leukemia and head and neck cancers (1, 2). FA is caused by inheritance of one of 22 dysfunctional FA genes (FANCA-FANCW) (3). Absence of any one member of the pathway causes genome instability during DNA replication, which results in mutagenic (cancer-causing) DNA damage and hypersensitivity to chemotherapeutic (normal and cancer-killing) DNA damage (4). Central to the FA pathway is the conjugation of ubiquitin to FANCI:FANCD2 (ID2) complexes (5, 6). ID2 monoubiquitination is critical to prevention of bone marrow failure in FA, but it is currently unknown how ID2-ub differs in its function to ID2. Several proteins have been proposed to specifically bind FANCI^Ub^ or FANCD2^Ub^ but not the un-ubiquitinated proteins (7, 8). For example, FAN1 nuclease was proposed to interact with FANCD2^Ub^ via its ubiquitin-binding domain (UBZ) (7), whereas recruitment of SLX4 endonuclease to the interstrand crosslink (ICL) site was shown to be dependent on FANCD2 ubiquitination (9). However, support for these interactions is limited to analysis of ubiquitination deficient (K>R) mutants, rather than evidence for direct ubiquitin-mediated protein interactions.

The retention of FANCD2 in chromatin foci is dependent on its monoubiquitination by a “core complex” of Fanconi anemia proteins (10). FANCI and the FA core complex are required to generate FANCD2-foci that mark the location of double strand breaks, stalled replication forks and R-loops (11-13) in the nucleus, and protect nascent DNA at these sites from degradation by cellular nucleases (14). The ubiquitinated form of FANCD2, and also its ubiquitinated partner protein FANCI, become resistant to detergent and high-salt extraction from these foci (15, 16), leading to speculation about the existence of a chromatin anchor or altered DNA binding specificity post-monoubiquitination (17).

A recent electron microscopy study revealed a DNA interacting domain that is required for FANCI:FANCD2 binding to DNA (18). The crystal structure of the non-ubiquitinated FANCI:FANCD2 shows that the monoubiquitination sites of FANCI:FANCD2 are buried and therefore inaccessible in the dimer interface of the complex (19), suggesting that DNA binding might be required to expose the ubiquitin binding sites. Based on biochemical analyses non-ubiquitinated FANCI and FANCD2 preferentially bind to branched DNA molecules which mimic DNA replication and repair intermediates (20-22), however how that activates monoubiquitination of FANCI:FANCD2 remains poorly understood. DNA is a cofactor for maximal ubiquitination (17, 23)

Here we have reconstituted the FA pathway using recombinant FA core complex and fluorescently labelled DNA oligomer substrates. We show that once monoubiquitinated, FANCI:FANCD2 forms a tight interaction with double-stranded containing DNA. We report the successful purification of monoubiquitinated FANCI:FANCD2 complex bound to DNA using an Avi-ubiquitin construct, and show that the monoubiquitination does not promote any new protein:protein interactions with other factors *in vitro*. Instead, we reveal a new role of monoubiquitinated FANCI:FANCD2 in forming higher order structures and demonstrate how monoubiquitinated FANCI:FANCD2 interacts with DNA to initiate DNA repair. Our work uncovers the molecular function of the pathogenetic defect in most cases of FA.

## Results

### Monoubiquitination does not promote association of FANCI:FANCD2 with a panel of proteins previously hypothesized to bind the ubiquitinated form

Mono-ubiquitinated FANCI:FANCD2 (henceforth I^Ub^D2^Ub^) is the active form of the complex in repair of DNA damage. Many previous studies have speculated about the existence of DNA repair proteins that specifically associate with I^Ub^D2^Ub^. A summary of these proteins is presented in Table 1. Using recombinant ID2 or I^Ub^D2^Ub^ prepared by a novel purification method (Figure 1a) (24) we sought to directly compare the binding of this panel of ID2-associated proteins. Each of the partner proteins was expressed using reticulocyte extracts (Figure 1b), and the majority bound to the ID2 complex as predicted based on previously identified associations (Figure 1c). The strongest binding proteins in terms of fraction of protein recovered were SLX4, SMARCAD, FANCJ, PSMD4, SF3B1, TRIM25, MCM5 and BRE. Luciferase protein was used as a control 35S-labeled prey-protein, and this protein did not bind to ID2 (Figure 1c). Surprisingly, we discovered that none of the proteins showed any increased affinity for I^Ub^D2^Ub^ over ID2 (Figure 1c-d).

**Figure 1.**
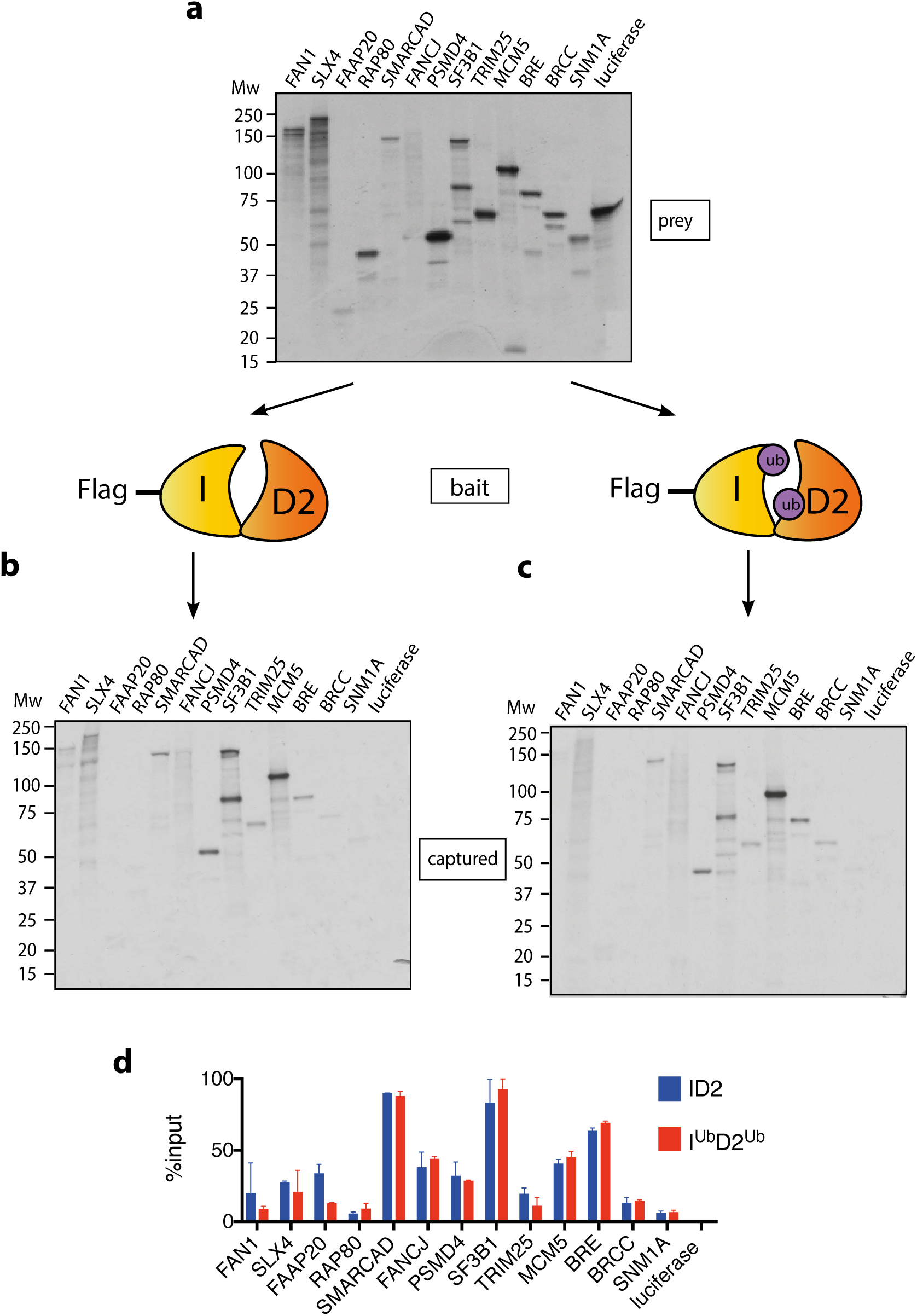
Mono-ubiquitination does not alter interaction of FANCI:FANCD2 with DNA repair proteins. (a) 35S-labelled FAN1, SLX4, FAAP20, RAP80, SMARCAD, FANCJ, PSMD4, SF3B1, TRIM25, MCM5, BRE, BRCC, SNM1A or luciferase (control) inputs were expressed using reticulocyte. (b) The inputs prepared from (c) were incubated with the indicated Flag-ID2 or (D) Flag-I^ub^D2^ub^, followed by Flag pull-down and elution. The complexes were subjected to SDS-PAGE, and radiolabeled proteins were detected by autoradiography (representative experiment of n=2). (d) Quantification showing percentage of ID2 or I^ub^D2^ub^ binding to inputs.

**Table 1:**
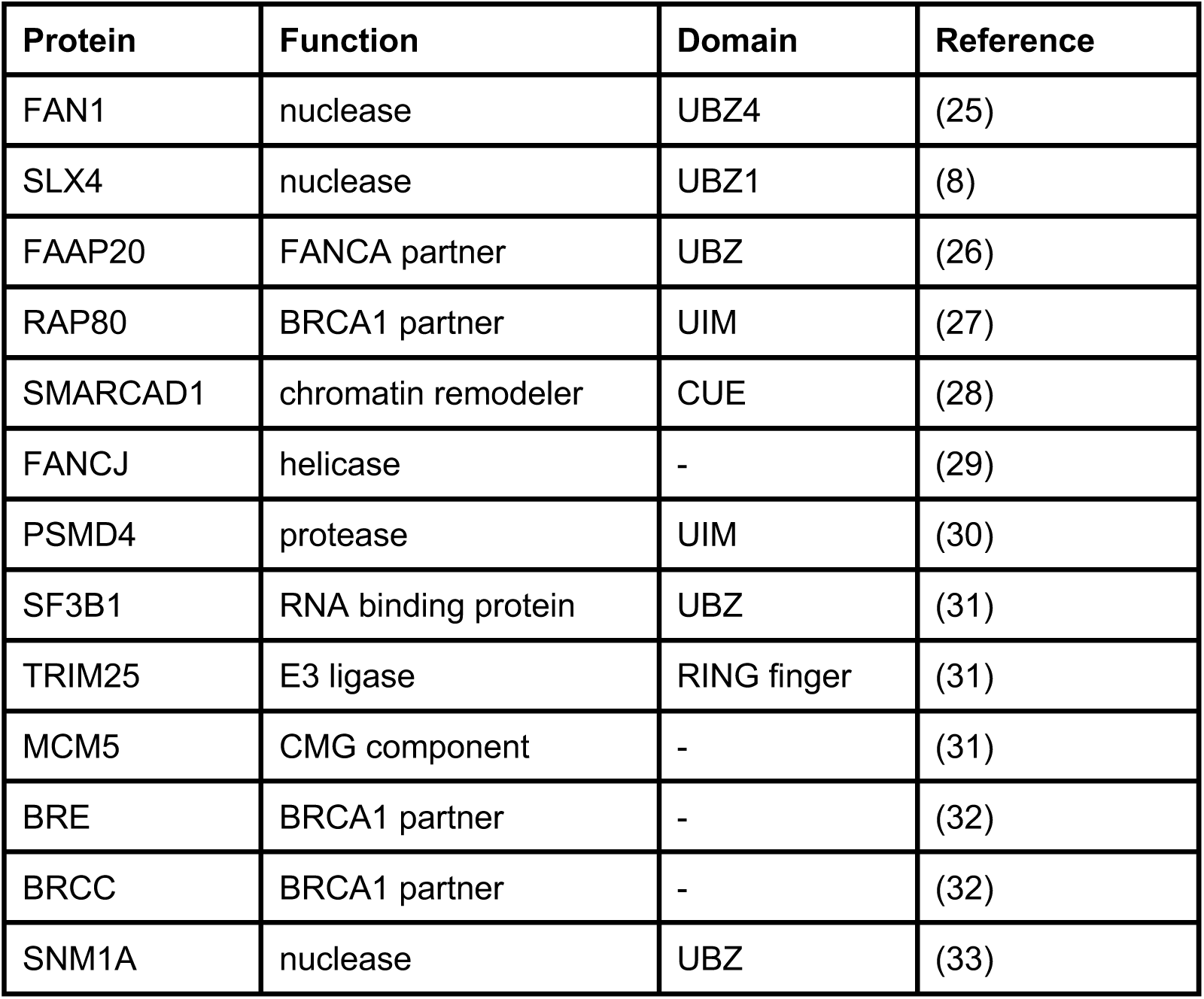
List of proteins containing ubiquitin binding domain that are described or predicted to bind to ubiquitinated FANCD2.

### Monoubiquitination locks FANCI:FANCD2 on DNA

An alternative explanation for the observed increased in association between ID2 and its associated proteins after DNA damage is that I^Ub^D2^Ub^ has an increased affinity for DNA, which brings the protein into closer proximity to these partners. The majority of ID2 associated proteins are chromatin localized. In order to explore the stability of I^Ub^D2^Ub^ on DNA, we performed *in vitro* monoubiquitination reactions in the presence of IR-dye700 labelled double-stranded DNA (dsDNA). As previously characterized (23), we observed DNA-dependent appearance of monoubiquitinated forms of FANCD2 and FANCI when using recombinant FA core complex components (Figure 2a-b). ID2 monoubiquitination readily lead to DNA mobility shifts using EMSA (electromobility shift assay) even at low concentrations, but this was not observed for the unmodified (apo)-ID2 complex in the absence of the enzymatically FA core complex, or when monoubiquination-defective K-to-R mutants of ID2 were used in the reaction (Figure 2c, lanes 2-4).

**Figure 2.**
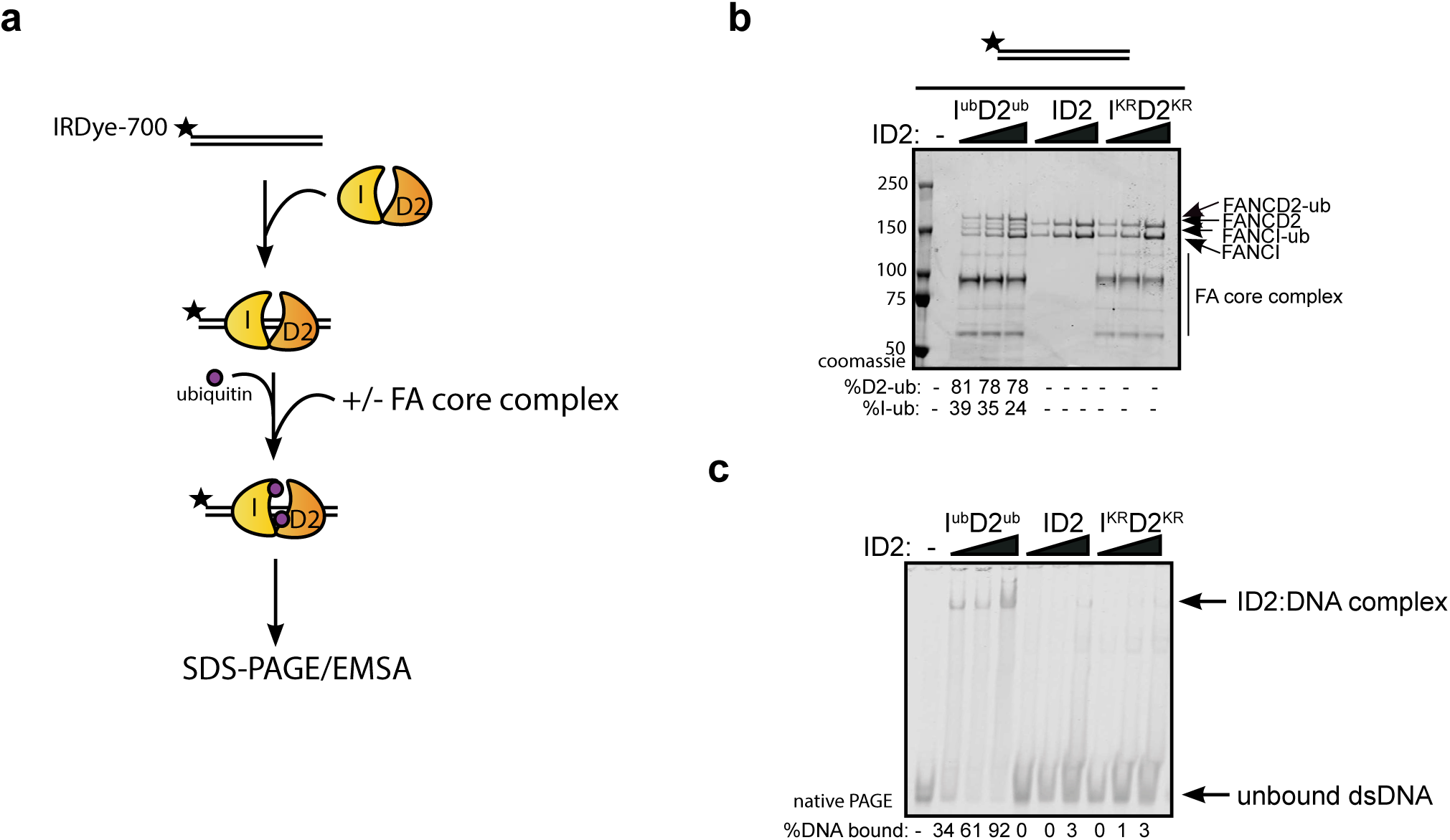
Monoubiquitination locks FANCI:FANCD2 on DNA. (A) Schematic of the electrophoretic mobility shift assay (EMSA) using IRDye-700 labelled dsDNA. (B) Coomassie stained SDS-PAGE gel showing monoubiquitination of FANCI:FANCD2 using recombinant FA core complex and IR-dye700 labelled dsDNA. 25, 50 and 100 nM of ID2 or I^KR^D2^KR^ were incubated with 25 nM of the IR-dye700 dsDNA for 90 min. The percentage of FANCI or FANCD2 monoubiquitination were calculated and shown under SDS-PAGE gel. (C) Monoubiquitination reactions from (B) were resolved on 6% native PAGE gel for EMSA analysis. The percentage of ID2 binding to DNA was calculated and shown under native PAGE gel.

Previously, we and others showed that various different dsDNA-containing structures could robustly stimulate ID2 monoubiquitination (20, 34, 35), but that single-stranded DNA (ssDNA) does not. To determine if monoubiquitinated ID2 had increased affinity for other dsDNA containing structures, we repeated the monoubiquitination reactions in the presence of different IR-dye700 labelled DNA structures. Each of the dsDNA containing structures, led to increased ID2 monoubiquitination and increased retention of an EMSA shifted band (Figure 3). Conversely, ssDNA very weakly stimulated monoubiquitination but did not cause an EMSA shift.

**Figure 3.**
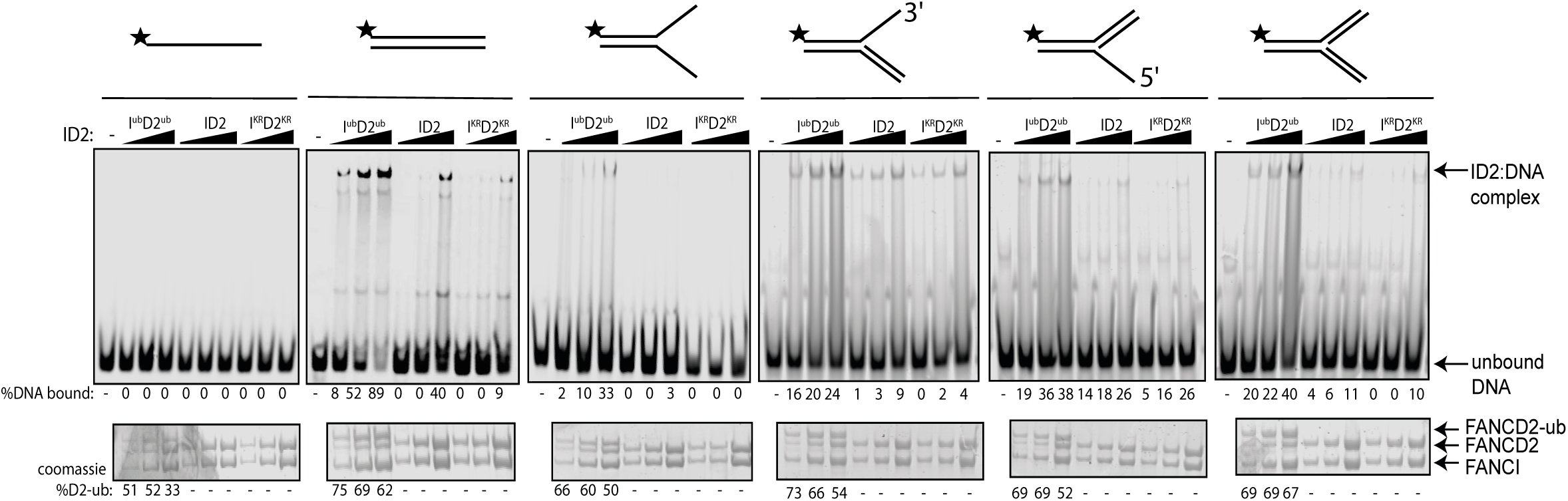
Monoubiquitinated FANCI:FANCD2 binds to any type of dsDNA. EMSA gels showing binding of monoubiquitinated or unmodified ID2 complex to different oligo-based DNA substrates. Above each panel, a schematic representing the tested DNA substrate is shown. 25, 50 and 100 nM of ID2 or I^KR^D2^KR^ were incubated with 25 nM of the indicated DNA substrate and the protein:DNA complexes were resolved on 6% PAGE gels (top). The percentage of DNA binding was calculated and shown under each EMSA gel. Coomassie stained SDS-PAGE gel (bottom) showing the ubiquitination reactions used in the EMSA. The percentage of FANCD2 ubiqitination was calculated and shown under each SDS-PAGE gel.

### Both FANCI^ub^ and FANCD2^ub^ are associated with a “locked” protein:DNA complex

Previous studies reported that monoubiquitination of ID2 complex may lead to dissociation of the heterodimer to its individual subunits, as measured by loss of co-immunoprecipitation of FANCI with FANCD2 (35, 36). In contrast, we did not observe any Ub-mediated dissociation of ID2 *in vitro*. First, Western blotting of the EMSA gels confirmed that the gel shifted DNA band contains both FANCI and FANCD2 proteins (Figure 4a). Second, FANCI^Ub^ still co-immunoprecipitated with FANCD2^Ub^ at the plateau of the in vitro ubiquitination reaction (Figure 4b).

**Figure 4.**
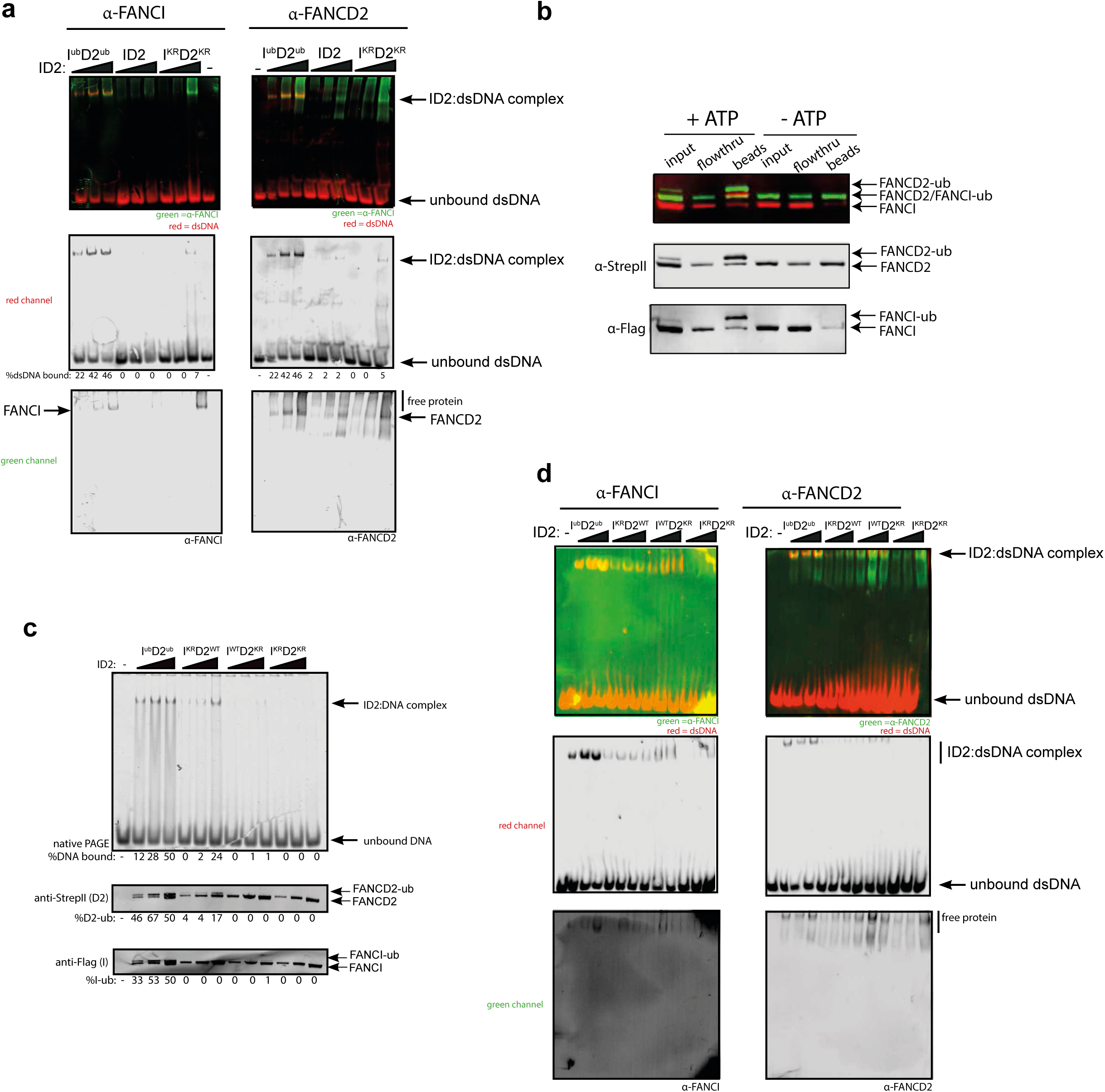
FANCD2 monoubiquitination is sufficient for FANCI:FANCD2 locking to DNA, but stimulated by FANCI monoubiquitination. (a) Western blots of the EMSA gels containing 25, 50 and 100 nM of I^ub^D2^ub^, ID2 or I^KR^D2^KR^ in the presence of 25 nM IRDye-700 labelled dsDNA (red). Left panels correspond to anti-FANCI antibody (green) and right panels correspond to anti-FANCD2 antibody (green) (b) StrepII affinity purification of mono-ubiquitinated (+ATP) and non-ubiquitinated ID2 (-ATP). (c) EMSA gels (top) and western blots (bottom) showing the monoubiquitination of 25, 50 and 100 nM I^WT^D2^WT^, I^KR^D2^WT^, I^WT^D2^KR^ or I^KR^D2^KR^ in the presence of 25 nM IRDye-700 labelled dsDNA. (d) Western blots of the EMSA gels from (C) showing FANCI (left, green) and FANCD2 (right, green) remained bound to IRDye-700 labelled DNA (red) after mono-ubiquitination.

To determine the contribution of each of FANCD2^Ub^ and FANCI^Ub^ to the locking of I^Ub^D2 ^Ub^ complex to DNA, we used ubiquitination-deficient (KR) mutants in the ubiquitination reaction. FANCI^KR^:FANCD2^WT^ or FANCI^WT^:FANCD2^KR^ mutant results in decrease in EMSA shift, and FANCI^KR^:FANCD2^KR^ did not bind to DNA (Figure 4c). However, this retention on DNA correlated with the extent of FANCD2 monoubiquitination retained by these mutant complexes. Western blotting the EMSA gels confirmed that both FANCD2 and FANCI are found in the EMSA shifted product, although in higher amounts when both proteins are capable of being monoubiquitinated (Figure 4d).

### Mutant forms of ubiquitin can still lock ID2 onto DNA

We postulated that the altered affinity for DNA induced by monoubiquitination must result from either a conformational change in the ID2 heterodimer after monoubiquitination, or participation of the conjugated ubiquitin directly in protein:DNA or protein:protein binding. To help distinguish these possibilities we utilized mutants of ubiquitin that have previously been shown to mediate the known protein:ubiquitin or protein:DNA interactions in other ubiquitinated protein interactions (Figure 5a) (37). Each of these Ub mutants were conjugated to ID2 by the FA core complex with similar efficiency (Figure 5b) and their locking onto DNA was then measured. Mutations in surface patch 1 (F4A, D58A), surface patch 2 (I44A, V70A), a DNA binding residue (K11R) or a tail mutant (L73P) had no apparent effect on DNA locking (Figure 5c). This result suggests that no canonical surface or region of ubiquitin is critical for DNA locking of ID2, and instead ubiquitin conjugation to ID2 probably induces a conformational rearrangement of the heterodimer.

**Figure 5.**
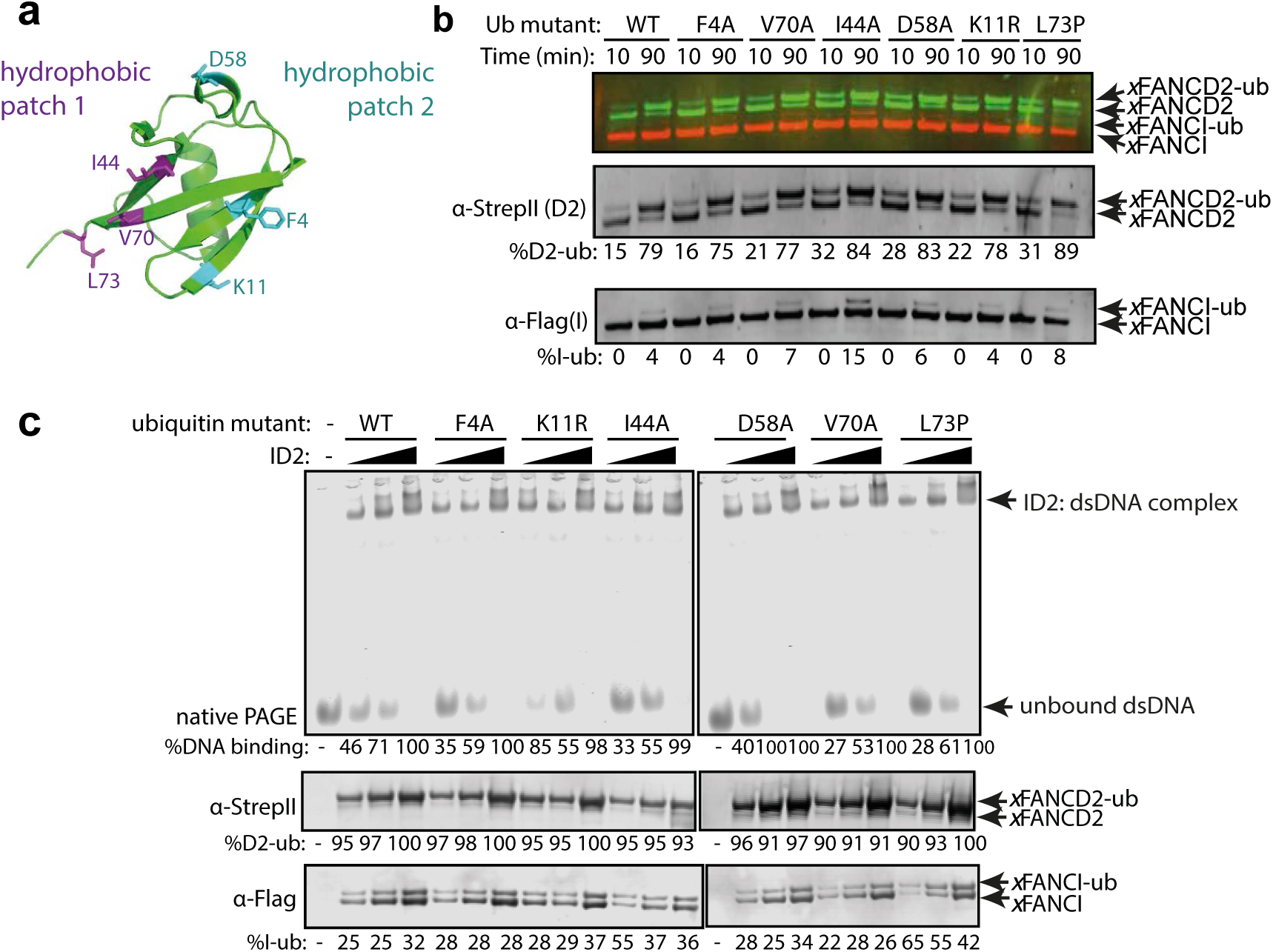
Mutation in different ubiquitin patches do not affect ID2 mono-ubiquitination or DNA binding. (a) Crystal structure of ubiquitin with ubiquitin mutant sites depicted (PDB: 1UBQ). Hydrophobic binding pockets are indicated in blue and pink. (b) Western blots showing the time course ubiquitination assays of ID2 using wild-type ubiquitin, ubiquitin mutant F4A, V70A, I44A, D58A, K11R and L73P. (c) EMSA gels showing 25, 50 and 100 nM monoubiquitinated ID2 binding to 25 nM IRDye-700 dsDNA using various ubiquitin mutants (top). Western blots of ID2 ubiquitination products were shown at the bottom and the percentage of FANCI and FANCD2 ubiquitination were shown at the bottom of each western blot panel.

### Purification of monoubiquitinated FANCI:FANCD2 complex bound to dsDNA reveals a filamentous architecture

In order to examine the architecture of purified recombinant I^Ub^D2^Ub^ complex in the presence of dsDNA plasmid, we utilized a recombinant Avi-tag ubiquitin construct containing a 3C protease site between the biotinylated Avi-tag and the N-terminus of ubiquitin (Figure 6a). This tagged ubiquitin is incorporated onto FANCI:FANCD2 by the FA core complex, allowing Avidin-Sepharose purification of monoubiquitinated ID2 that is then eluted by 3C protease cleavage. We recovered monoubiquitinated FANCI:FANCD2 complex only when FANCI is monoubiquitinated, suggesting that the N-terminus of D2-attached ubiquitin may be buried within the di-ubiquitinated complex, but the N-terminus of ubiquitin attached to FANCI is accessible for streptavidin binding (Figure 6b).

**Figure 6.**
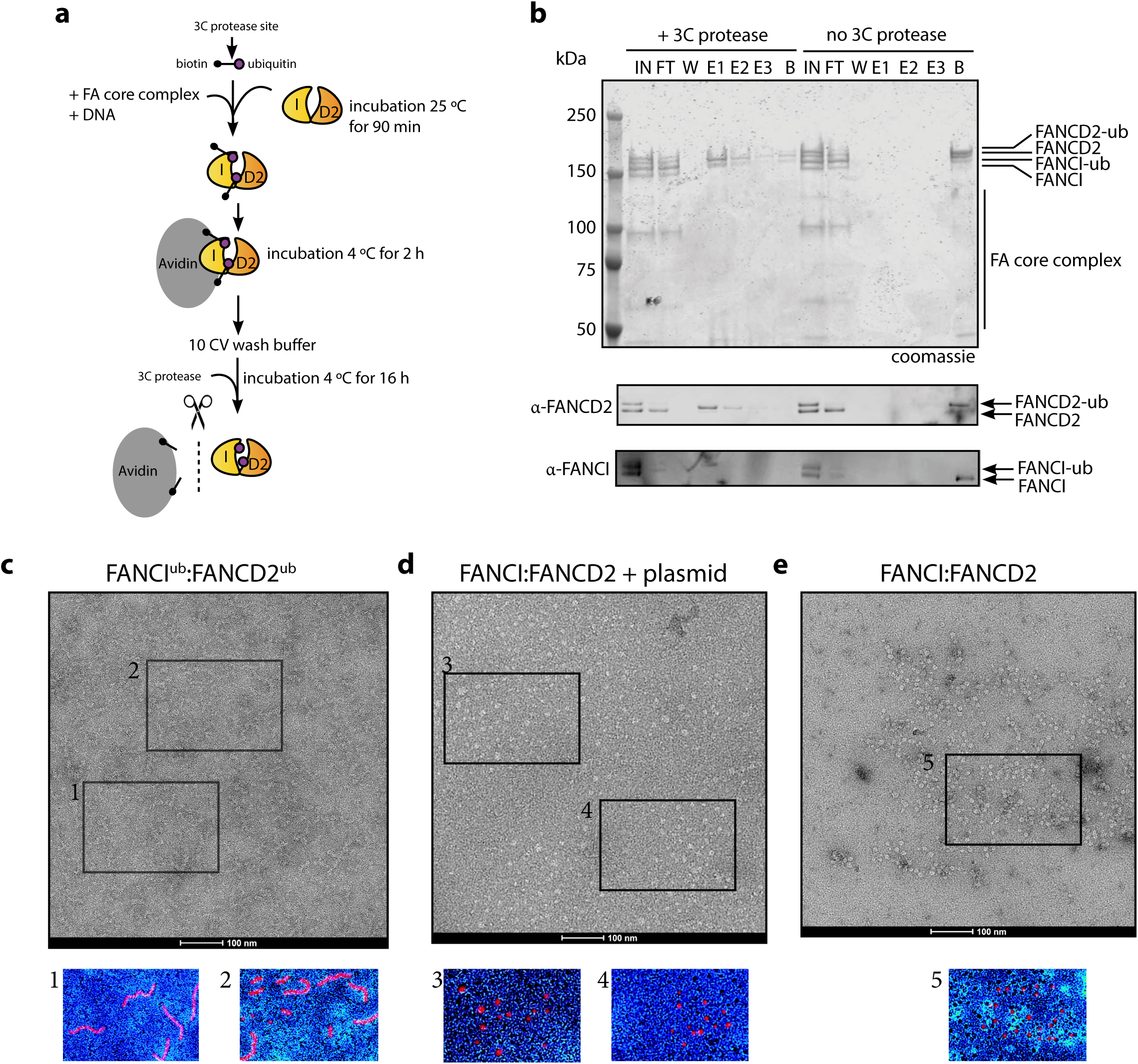
Mono-ubiquitinated FANCI:FANCD2 complex assemble into filament-like arrays. (a) Schematic of purification of monoubiquitinated FANCI:FANCD2 using Avi-ubiquitin. (b) Coomassie stained SDS-PAGE gel (top) and western blots (bottom) showing the purification of monoubiquitinated FANCI:FANCD2 complex eluted using Prescission protease (lanes 1-7) compared to without Prescission protease (lanes 8-14). (c-e) Representative negative-stained EM image of purified FANCI^ub^:FANCD2^ub^ complex bound to 2.7 kb plasmid, unmodified FANCI:FANCD2 incubated with 2.7 kb plasmid and unmodified FANCI:FANCD2 complex. Pseudo-coloured regions are shown to highlight particular filament-like arrays in FANCI^ub^:FANCD2^ub^ but not other samples.

Using this purified protein, we compared FANCI^ub^:FANCD2^ub^ to unmodified FANCI:FANCD2 using electron microscopy (EM). Surprisingly, we observed that FANCI^ub^:FANCD2^ub^ forms filament-like oligomers when bound to dsDNA plasmid (Figure 6c). Such filaments were not observed in the unmodified FANCI:FANCD2 protein preparation in the absence of presence of plasmid DNA, nor in previous investigations of human or Xenopus FANCI:FANCD2 complexes studied by EM (Figure 6d-e and (38-40)).

When smaller DNA molecules were used as the substrate for ID2 binding, we either observed no filament-like structures (60bp, Figure 7a) or shorter filament-like structures (150bp, Figure 7b) compared to structures that were on average 7-8x longer than the characteristic double saxophone structure of ID2 heterodimer in the non-ubiquitinated state (Figure 7c).

**Figure 7.**
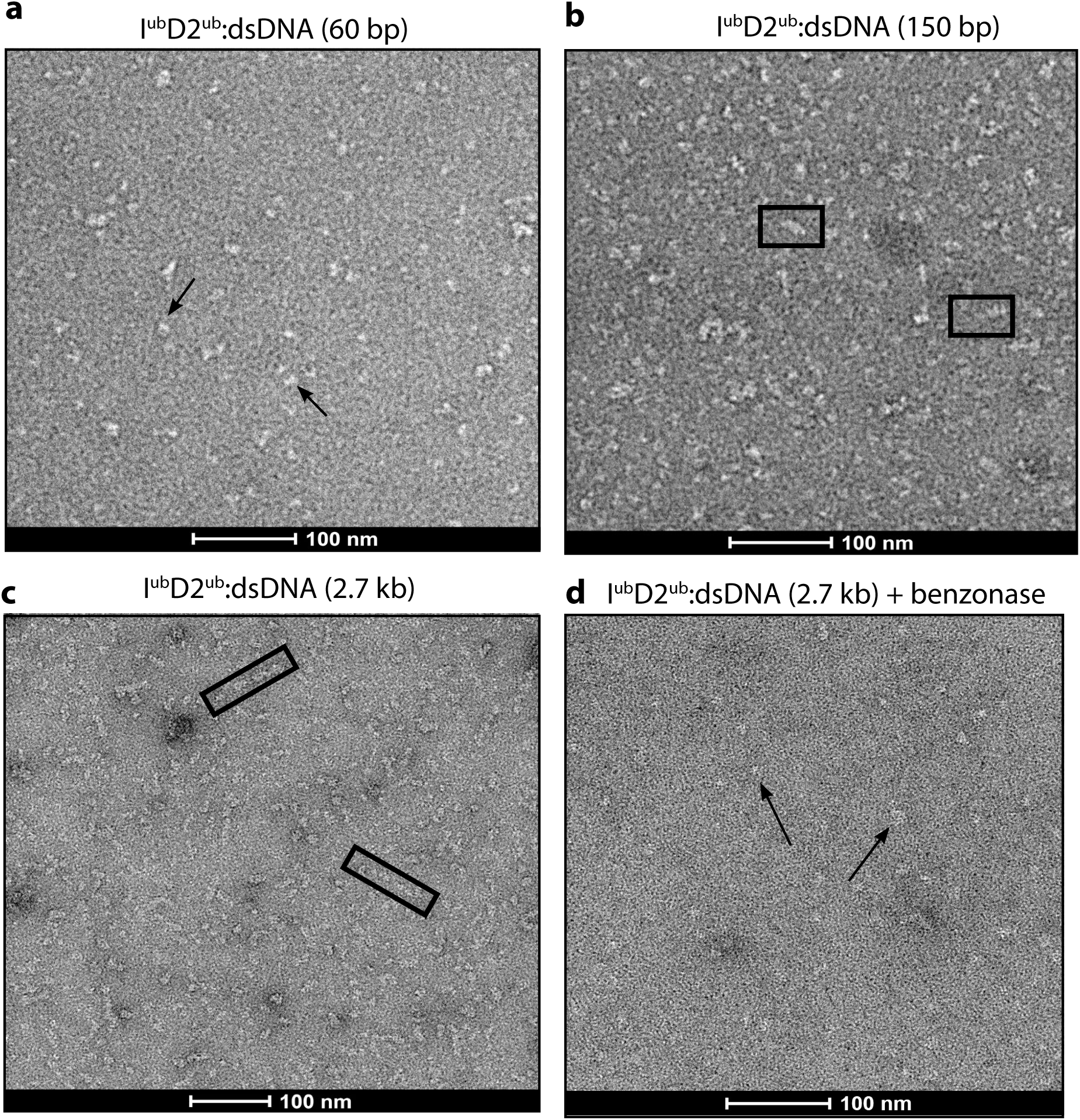
Monoubiquitinated FANCI:FANCD2 forms filamentous oligomers along the length of dsDNA. (A-D) Representative EM image of monoubiquitinated FANCI:FANCD2 bound to (A) 60 bp dsDNA, (B) 150 bp dsDNA, (C) kb dsDNA, and (D) 2.7 kb dsDNA and benzonase-treated.

The observation that filament length correlated with the size of DNA available for ID2 binding strongly suggested that the association between heterodimer subunits in the array was DNA-mediated. To test whether the array of I^ub^D2^ub^ is also dependent upon binding to the same DNA molecule we examined the plasmid-stimulated ubiquitination reaction products after treatment with the non-specific endonuclease, Benzonase. It is apparent from EM images that addition of Benzonase breaks the long filaments formed by I^ub^D2^ub^ complex into very short or heterodimer-sized units (Figure 7d). This finding is consistent with Benzonase cleaving exposed DNA between I^ub^D2^ub^ units, leading to destabilization of the filamentous arrays. Together our results show that, *in vitro*, ubiquitination of ID2 leads to a ubiquitin- and DNA-stabilized filamentous structure.

### Single I^ub^D2^ub^ heterodimers on short 60bp DNA have an altered architecture

Due to variability in the length and shape of filament-like I^ub^D2^ub^ structures on longer DNA molecules we have not been able to uncover the shape or subunit rearrangement of the individual units of the arrays. However, examination of I^ub^D2^ub^ purified together with short 60bp DNA allowed us to collect sufficient images of individual particles for analysis. These particles were similar in size to non-ubiquitinated ID2, but it is clear from individual molecule and class average views that the I^ub^D2^ub^ complex forms a distinct architecture from that of ID2 (Figure 8). In particular, the overall shape of individual particles and their class averages reveal a twisting that repositions the solenoid arms of one or both of the subunits bringing them into closer proximity. The conformational change induced appears to reduce the size of ID2 in an X but not Y direction, similar to that predicted in a previously model prediction that placed DNA in a channel between FANCI and FANCD2 post DNA binding (17). These images support the view that monoubiquitination induces a conformational change in the ID2 complex that locks it upon DNA.

**Figure 8.**
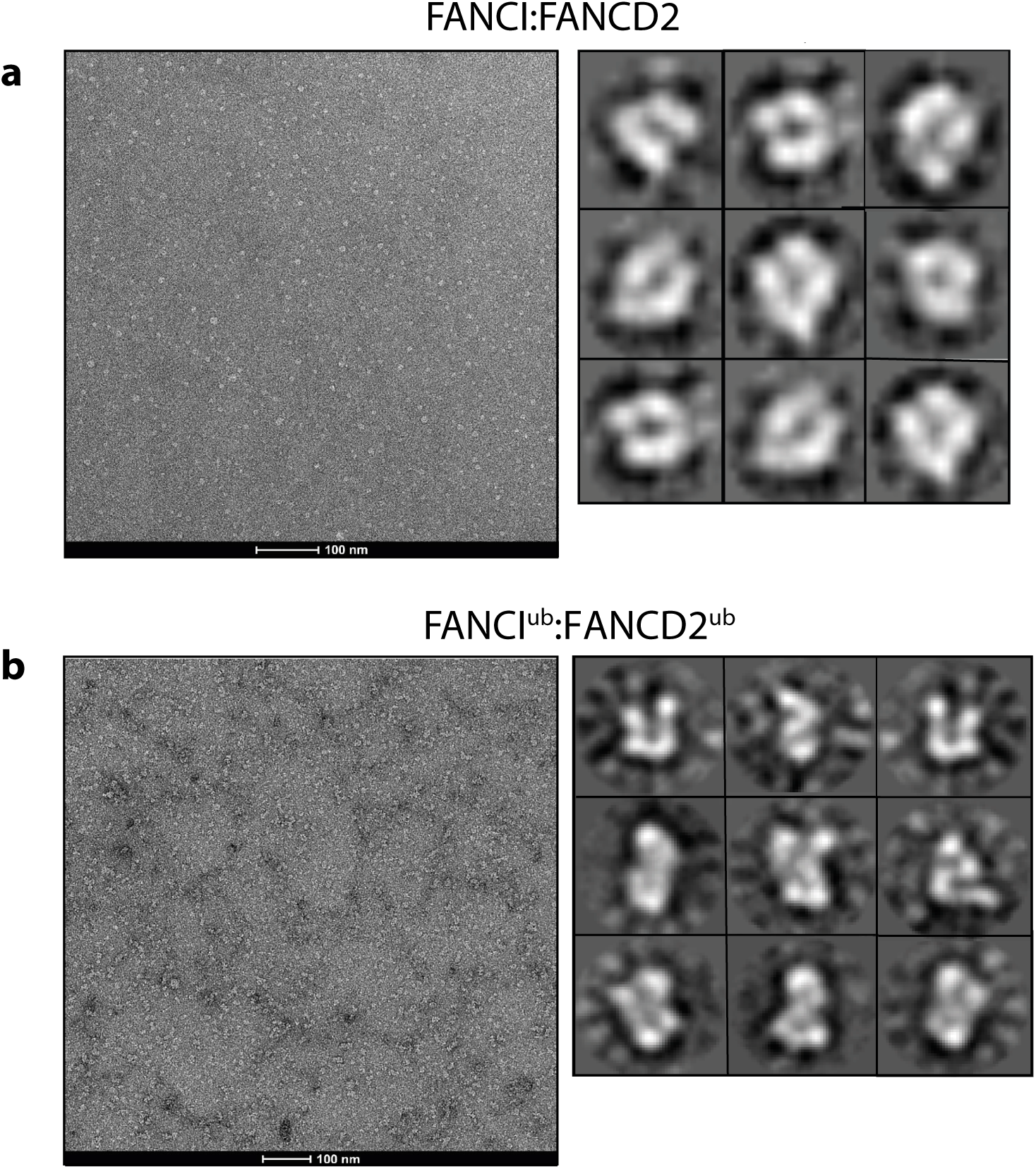
Electron micrographs of ID2 and I^ub^D2^ub^. (A) Representative negative-stained electron micrograph and 2D class average of ID2. (B) Representative negative-stain electron micrograph and 2D class average of I^ub^D2^ub^.

## Discussion

The protection of stalled forks by DNA repair factors is essential for proper DNA replication and the maintenance of genome stability. The primary mechanism of replication fork stabilization at ICLs, and other blocking damage, utilizes the proteins of the FA-BRCA DNA repair pathway. Monoubiquitination of FANCI:FANCD2 by the FA core complex is the central event in this pathway. Here we showed that monoubiquitination directly locks the ID2 complex onto double-stranded DNA, and promotes a filament-like coating of long DNA molecules. This finding answers a long-standing question about the nature of the biochemical reaction that is absent in the majority of patients with FA (41, 42).

“Fork protection” involves (i) exclusion of cellular nucleases such as MRE11 and DNA2 from the stalled DNA replication fork, and (ii) specific recruitment of other factors that are able to restart DNA replication (14, 43, 44). The role of nuclease exclusion seems most important because inhibition of either of two nucleases, MRE11 or DNA2, can significantly alleviate the sensitivity of FA-BRCA mutant cells to replication stalling agents (14, 43). But in many studies, specific association of various repair factors with I^Ub^D2^Ub^ compared to ID2 have been described. These proteins mostly contain ubiquitin-binding domains, providing an impetus for the recruitment-based hypothesis (summary and references in Table 1). However, because these previous studies focused on use of ubiquitination-deficient mutants, they could not address the underlying question of whether ubiquitin on either FANCI, FANCD2 or both proteins directly mediated the interaction. Now, we have shown that none of these proteins specifically bind recombinant purified I^Ub^D2^Ub^ compared to ID2. This finding was unexpected, and included proteins such as SLX4, MCM5, and FAN1, which have a demonstrated and essential role in the FA-pathway (10, 31, 45).

Instead, our data provide direct evidence that ID2 undergoes a conformational change after monoubiquitination, that leads to it becoming locked on double-stranded DNA. It is likely that in previous cell-based experiments specific interaction between SLX4 and FAN1 with FANCD2 but not FANCD2^K561R^ was observed because FANCD2^K561R^ does not become locked on DNA, even in the presence of active FA core complex. This finding is supported by the observation that FANCD2^K561R^ also does not becomes chromatin localized at damage sites even after extensive DNA damage (ref). We propose that I^Ub^D2^Ub^ does not demonstrate restricted interaction with any specific protein partner. None-the-less, its retention in chromatin after ubiquitination would be much more likely to bring the complex into proximity of these other DNA repair factors, where it could still influence their activity.

Locking onto DNA occurs through a ubiquitin-mediated conformational change in the ID2 complex. A ubiquitin binding-domain (UBD) in FANCD2 has previously been shown necessary for the retention of the protein in the chromatin faction, and for strong binding to FANCI (46). This UBD domain sits in the FANCD2 structure opposite to where ubiquitin is likely to reside after its conjugation on to FANCI by the FA core complex, and most likely mediates the locking function and conformational rearrangement. Relocation of a FANCD2 tower domain upon DNA binding most likely stimulates the conformational rearrangement that is then stapled in place by ubiquitin:UBD association. Other DNA binding proteins such as histone H2A show an increased association with DNA after monoubiquitination (47) and monoubiquitination also increases the DNA occupancy of transcription factors such as FOXO4 and CIITA (48). It’s possible that ubiquitin to UBD mediated locking is a general mechanism of protein:DNA target stabilization.

### I^Ub^D2^Ub^ locked in nucleoprotein filaments

In addition to a conformational change in ID2 induced by monoubiquitination (that has also been concurrently discovered and reported by the Paveltich and Passmore labs (49, 50)) we found that monoubiquitinated I^Ub^D2^Ub^ formed large filament-like arrays when it was purified together with plasmid DNA, but not short 60bp DNA fragments. On average, the length of plasmid-associated structures is 7-8x that of that associated with 60bp DNA. Larger or longer arrays may potentially be obscured from view because the purification strategy makes elution exponentially more difficult with increasing numbers of conjugated ubiquitin-molecules. Steps to remove “aggregates” may have also inadvertently removed larger arrays. However, as the number of potential plasmid DNA binding sites for ID2 was in large excess the concentration of ID2 used to stimulate reaction, their appears to be some purpose to creation of these filamentous arrays.

There is evidence that I^Ub^D2^Ub^ locked in nucleoprotein filaments exist in cells. Antibodies against FANCD2 have long been used as a marker of double strand breaks, stalled replication forks and R-loops because the protein forms large, intensely staining foci during S-phase that are increased after treatment with DNA damaging agents (11, 12, 51). We suspect that these intense foci are due to coating of DNA around damaged forks, potentially in filamentous arrays similar to those we observed by EM. Support for the large size and extent of DNA binding reflective of filamentous arrays also comes from chromatin immunoprecipitation and sequencing (ChIP-Seq) using anti-FANCD2 (13). FANCD2, and two other damage markers MRE11 and γH2AX, showed no specific localization in a bulk population of cells, but strongly localized adjacent to a Cas9-induced site-specific DNA break. Both γH2AX and FANCD2 produced a broad peak centred at the target site kilobases (kb) to megabases (mb) in length. In contrast, MRE11 is located within a very tight peak within ∼100bp of the break. Chromatin within 1-2 kb of the DSB showed reduced occupancy by γH2AX, consistent with dechromatinization around break sites (52), but FANCD2 was present right up to the DSB. Accumulation of FANCD2 increases at the DSB early after cleavage, and accumulates more distant from the DSB progressively with time post-cleavage. This is suggestive of a polymerisation of the FANCD2 signal away from the break site, as hypothesised would occur for a protein that forms a growing filament at broken DNA (13). The conserved function of FANCD2 as a histone chaperone (53, 54) may even be directly linked to displacement of nucleosomes as filamentous arrays extend into break-adjacent chromatin.

In this study, we also observed direct association of two ID2 heterodimers by co-immunopurification only after the protein becomes monoubiquitinated. This approach, if performed in cells, could be used to further delineate the mechanism and cellular factors required for the extension of I^Ub^D2^Ub^ arrays during fork protection. Of particular interest will be determining the role of BRCA1 in locking and/or array extension. BRCA1:BARD1 was initially thought to be the E3 for FANCD2 monoubiquitination, because it co-immunoprecipitates FANCD2, and FANCD2 does not form nuclear foci after damage in BRCA1-deficient cells [31]. However, in various assays it was later shown that FANCD2 monoubiquitination does occur in BRCA1-deficient cells, but it is *uncoupled* from FANCD2 foci formation [32, 33].

### How would a locked ID2 filament mediate fork protection?

Filamentous structures on DNA play a genome protective role in prokaryotes: eg DAN protein forms a rigid collaborative filament that reduces accessibility during anoxia (55), while the *Vibrio cholera* protein ParA2 forms protective filamentous structures on DNA during segregation (56). Structural characterization has demonstrated how these filaments function and, in the case of ParA2, can be targeted therapeutically (57). The coating of ssDNA by RPA in eukaryotes, also protect DNA from the activity of nucleases, and directs the specific activity of others (58-60). We propose that a FANCI^ub^:D2^ub^ filament may have a similar stabilising role on newly synthesised dsDNA at a stalled replication fork. This property would explain why stalled forks are prone to degradation in FA and BRCA patient cells (14, 43). In particular, we hypothesise that filamentous DNA-locked I^ub^:D2^ub^ could prevent access to DNA by MRE11 and DNA2 nucleases and prevent aberrant ligation of broken DNA to other parts of the genome by non-homologous end-joining.

Second, the tight binding of FANCI^ub^:D2^ub^ to dsDNA, when localized to stalled replication forks, may also prevent the branch migration of replication forks and prevent their spontaneous or helicase-mediated reversal (61). Reversed forks are the substrate for degradation by DNA2 and WRN nuclease activities, providing a hypothetical link between the activities of FANCD2-monoubiquitination and the nuclease activity of DNA2 and WRN (62, 63)

Third, I^ub^:D2^ub^ arrays may also locally suppress non-homologous end-joining (NHEJ) factors, and/or delineate the newly synthesized chromatin from unreplicated regions during the promotion of templated repair processes such as homologous recombination. FANCD2, FANCI, and components of the FA core complex were identified amongst relatively few other factors, in a genome-wide screen for genes that promote templated repair over NHEJ (64). Stabilization of RAD51 filaments, required for HR, is also an *in vitro* property of ID2 (65), suggesting I^ub^:D2^ub^ filamentous arrays may exist adjacent to or coincident with RAD51 filaments in cells, in order to provide a polarity to the homologous recombination reaction without loss or gain of genomic sequences.

### Role of FANCI-monoubiquitination

Fanci and Fancd2 have common and distinct functions in mouse models of Fanconi anemia (66), while the double knockout of FANCI and FANCD2 has an unexpectedly distinct phenotype compared to single knockouts in human cells (67). But FANCI^K523R^ expressing cells are less sensitive to DNA damage than FANCI knockout in human cells(15), so what is the role of FANCI monoubiquitination? Previous studies demonstrated that FANCI monoubiquitination is always subsequent to FANCD2 monoubiquitination, both in cells (68) and in biochemical assays (23). FANCI also likely plays a role in recruiting the FA core complex to the substrate (69). In this study, we show that FANCI-monoubiquitination is not necessary for locking of the ID2 complex onto DNA (Figure 4). However, *in vivo* it is likely that FANCI monoubiquitination plays a critical role in regulating deubiquitination of the ID2 complex. FANCI recruits the deubiquitinating enzyme USP1:UAF1 (70), which prevents trapping of monoubiquitinated FANCD2 at non-productive DNA damage sites, but only ID2^Ub^ but not I^Ub^D2^Ub^ is a substrate (23). It is also clear from our EM investigations that FANCI must play an important role in the structural integrity of I^Ub^D2^Ub^ filamentous arrays on DNA, possibly creating an asymmetry necessary for a specific polarity to array assembly.

### Implications for understanding the deficiency of Fanconi anemia

Onset of progressive bone marrow failure occurs at a median age of 7 in children with FA (71). Almost all of these patients lack FANCD2 and FANCI monoubiquitination, due to mutation in either *FANCD2* or *FANCI* or one of the 9 other FANC proteins required for their monoubiquitination (10). The importance of the monoubiquitin signal is highlighted by the observation that up to 20% of patients acquire somatic reversion of the inherited mutation in a fraction of blood cells (72). These mutations restore monoubiquitination and prevent bone marrow failure. Our work suggests two potential strategies for treatment of FA: restoration of gene function, such as that which occurs in somatic revertants or, identification of novel mechanisms to stabilize an ID2:DNA-locked complex for fork protection by ubiquitin-mediated or innovative means. New small molecule activators or inhibitors of ID2:DNA locking could be therapeutics in FA or cancer-treatment. *In vitro* biochemistry has proven to be the most powerful tool in uncovering new functions of FANCD2-monoubiquination that had gone undiscovered for nearly 20 years. The approach is likely to be formidable in drugging the FA pathway in future studies.

## Materials and Methods

### Protein purification

Flag-FANCI and StrepII-FANCD2 were expressed using the pFastBac1 vector (Life Technologies). For FANCI:FANCD2 complex, Hi5 cell pellets were resuspended in lysis buffer (50 mM Tris-HCl pH 8.0, 0.1 M NaCl, 1 mM EDTA, 10% glycerol and 1X mammalian protease inhibitor), and sonicated. Lysates were clarified by centrifugation at 20,000g and the supernatants were incubated with M2 anti-FLAG agarose resin for 2 h. The resin was washed 5×5 min incubation with wash buffer (20 mM Tris-HCl pH 8.0, 0.1 M NaCl, 10% glycerol), and the protein was eluted in the same buffer containing 100 μg/mL FLAG peptide. GST-UBE2T, Flag-BL100, MBP-CEF were purified as described (Van Twest, 2017). His-UBE1 was purchased from Boston Biochem.

### Biotinylated-Avi-ubiquitin purification

His-Avi-ubiquitin was purified as described in (24).

### *In vitro* ubiquitination assay

Standard ubiquitination reactions contained 10 μM recombinant human avidin-biotin-ubiquitin, 50 nM human recombinant UBE1, 100 nM UBE2T, 100 nM PUC19 plasmid, 2 mM ATP, 100 nM FANCI:FANCD2 complex wild type (WT) or ubiquitination-deficient (KR), in reaction buffer (50 mM Tris-HCl pH 7.4, 2.5 mM MgCl_2_, 150 mM NaCl, 0.01% Triton X-100). 20 μL reactions were set up on ice and incubated at 25°C for 90 minutes. Reactions were stopped by adding 10 μL NuPage LDS sample buffer and heated at 80°C for 5 minutes. Reactions were loaded onto 4-12% SDS PAGE and run using NuPAGE ® MOPS buffer and assessed by western blot analysis using Flag (Jomar Life Research) or StrepII (Abcam) antibody.

### *In vitro* Transcription/translation pull down of ^35^S-labeled proteins

Flag-tagged FANCI:FANCD2 and monoubiquitinated FANCI:FANCD2 was prepared by incubating purified FANCI:FANCD2 or monoubiquitinated FANCI:FANCD2 on Flag beads for 2 hr followed by extensive washes in buffer A (20 mM TEA pH 8.0, 150 mM NaCl, 10% glycerol). ^35^S-labeled proteins containing UBZ or other ubiquitin domains (Table 1) were generated using the TNT Quick Coupled T7 Transcription/Translation System (Promega) and ^35^S-labeled methionine (Perkin Elmer). 10 μL of TNT product was incubated for 4 hr at 4 °C in buffer A with 100 ng Flag-tagged FANCI:FANCD2 or monoubiquitinated FANCI:FANCD2, 20 μL of Flag-beads (Sigma-Aldrich) in a 100 μL reaction. Beads were washed five times with buffer A and resuspended in LDS loading buffer. Proteins were separated by SDS-PAGE and visualized by autoradiography.

### Electrophoretic mobility shift assay

Oligonucleotides used to create fluorescently labeled DNA were IRDYE-700-labelled X0m1 (IDTDNA) and other oligos with the sequences shown in Supplementary Table 1. Assembly of the different DNA structures was performed exactly as previously described (Supplementary Table 2, (23)). 25 nM DNA substrates were incubated in 20 μL ubiquitination buffer containing 100 nM FANCI:FANCD2, 100 nM BL100, 100 nM CEF, 10 uM HA-ubiquitin (Boston Biochem), 50 nM UBE1 (Boston Biochem) and 100 nM UBE2T at room temperature for 90 min to initiate ubiquitination. The reaction was resolved by electrophoresis through a 6% non-denaturing polyacrylamide gel in TBE (100 mM Tris, 90 mM boric acid, 1 mM EDTA) buffer and visualized by Licor Odyssey system. Competitive EMSA were performed by adding cold dsDNA at concentrations 0, 0, 25, 2.5, 7.5, 25, 75, 250 and 1250 nM during or after the ubiquitination reaction.

### Purification of monoubiquitinated FANCI:FANCD2 complex

Di-monoubiquitinated FANCI:FANCD2 complex was purified as described (24). DNA molecules of 60bp or 150bp (dsDNA from oligonucleotides) or 2.6kb (circular plasmid DNA) were used to stimulate the reaction for different experiments, as indicated.

### Mass spectrometry analysis of monoubiquitinated FANCI:FANCD2 complex

Gels containing monoubiquitinated FANCI and FANCD2 bands were excised and in-gel digested with trypsin and subjected to LC/MS analysis on ESI-FTICR mass spectrometer at Bio21. The analysis program MASCOT was used to identify ubiquitination sites on FANCI and FANCD2.

### Negative stained electron microscopy

Freshly purified monoubiquitinated or non-ubiquitinated FANCI:FANCD2 complex was applied to glow-discharged, carbon/formvar grids and allowed to adsorb for 60 s. Specimen was then stained with 2% uranyl formate for 60 s. Specimen were imaged at a magnification of 73,000 x with camera (corresponding to a pixel size of 1.9 A) in Tecnai 120 kV.

### Single-particle image processing

Monoubiquitinated or non-ubiquitinated FANCI:FANCD2 particles were semi-automatically picked using XMIPP3 (73). The parameters of the contrast transfer function (CTF) for negative stained data was estimated on each micrograph using CTFFIND3 (74). Finally, reference free 2D alignment and averaging were executed using XMIPP3 or CisTEM (75).

## Acknowledgements

We thank Alexandra Sobeck, Puck Knipscheer, Steve West, Johan de Winter, KJ Patel, Paul Hasty, Dario Alessi, Timothy Richmond, Beverlee Buzon, Andrew Blackford, Steve Jackson and Stephen Elledge for reagents. We thank Eric Hanssen from the Electron Microscopy facility, and Nick Shuai from the Mass Spectrometry facility, at Bio21 Institute, Melbourne. WT was supported by an Australian Government Research Training Scheme postgraduate scholarship. AJD is a Victorian Cancer Agency fellow. WMC is an NHMRC career development fellow and Maddie Riewoldt’s vision fellow (WC-MRV2016). MWP is an NHMRC Australia Senior Research Fellow. This work was funded by grants from the Fanconi Anemia Research Fund (to AJD and WC), Maddie Riewoldt’s Vision (to WC), the National Health and Medical Research Council (GNT1123100 and GNT1181110 to AJD and GNT1156343 to WMC), and the Victorian Government’s OIS Program.

## References

1. Gillio AP, Verlander PC, Batish SD, Giampietro PF, Auerbach AD. Phenotypic Consequences of Mutations in the Fanconi Anemia &lt;em&gt;FAC&lt;/em&gt; Gene: An International Fanconi Anemia Registry Study. Blood. 1997;90(1):105

2. Butturini A, Gale RP, Verlander PC, Adler-Brecher B, Gillio AP, Auerbach AD. Hematologic abnormalities in Fanconi anemia: an International Fanconi Anemia Registry study. Blood. 1994;84(5):1650

3. Tan W, Deans AJ. A defined role for multiple Fanconi anemia gene products in DNA-damage-associated ubiquitination. Experimental Hematology. 2017;50:27–32 https://doi.org/10.1016/j.exphem.2017.03.001.

4. Deans AJ, West SC. DNA interstrand crosslink repair and cancer. Nature reviews Cancer. 2011;11(7):467–80 https://doi.org/10.1038/nrc3088.

5. Walden H, Deans AJ. The Fanconi Anemia DNA Repair Pathway: Structural and Functional Insights into a Complex Disorder. Annual Review of Biophysics. 2014;43(1):257–78 https://doi.org/10.1146/annurev-biophys-051013-022737.

6. Chirnomas D, Taniguchi T, de la Vega M, Vaidya AP, Vasserman M, Hartman A-R, et al. Chemosensitization to cisplatin by inhibitors of the Fanconi anemia/BRCA pathway. Molecular Cancer Therapeutics. 2006;5(4):952 https://doi.org/10.1158/1535-7163.MCT-05-0493.

7. Smogorzewska A, Desetty R, Saito TT, Schlabach M, Lach FP, Sowa ME, et al. A genetic screen identifies FAN1, a Fanconi anemia associated nuclease necessary for DNA interstrand crosslink repair. Molecular cell. 2010;39(1):36–47 https://doi.org/10.1016/j.molcel.2010.06.023.

8. Lachaud C, Castor D, Hain K, Muñoz I, Wilson J, MacArtney TJ, et al. Distinct functional roles for the two SLX4 ubiquitin-binding UBZ domains mutated in Fanconi anemia. J Cell Sci. 2014;127(13):2811

9. Klein Douwel D, Boonen Rick ACM, Long David T, Szypowska Anna A, Räschle M, Walter Johannes C, et al. XPF-ERCC1 Acts in Unhooking DNA Interstrand Crosslinks in Cooperation with FANCD2 and FANCP/SLX4. Molecular Cell. 2014;54(3):460–71 https://doi.org/10.1016/j.molcel.2014.03.015.

10. Walden H, Deans AJ. The Fanconi anemia DNA repair pathway: structural and functional insights into a complex disorder. Annual review of biophysics. 2014;43:257–78 https://doi.org/10.1146/annurev-biophys-051013-022737.

11. Taniguchi T, Garcia-Higuera I, Andreassen PR, Gregory RC, Grompe M, D’Andrea AD. S-phase-specific interaction of the Fanconi anemia protein, FANCD2, with BRCA1 and RAD51. Blood. 2002;100(7):2414–20

12. Schwab RA, Nieminuszczy J, Shah F, Langton J, Lopez Martinez D, Liang CC, et al. The Fanconi Anemia Pathway Maintains Genome Stability by Coordinating Replication and Transcription. Mol Cell. 2015;60(3):351–61 https://doi.org/10.1016/j.molcel.2015.09.012.

13. Wienert B, Wyman SK, Richardson CD, Yeh CD, Akcakaya P, Porritt MJ, et al. Unbiased detection of CRISPR off-targets in vivo using DISCOVER-Seq. Science. 2019;364(6437):286–9 https://doi.org/10.1126/science.aav9023.

14. Schlacher K, Wu H, Jasin M. A distinct replication fork protection pathway connects Fanconi anemia tumor suppressors to RAD51-BRCA1/2. Cancer Cell. 2012;22(1):106–16 https://doi.org/10.1016/j.ccr.2012.05.015.

15. Smogorzewska A, Matsuoka S, Vinciguerra P, McDonald ER, Hurov KE, Luo J, et al. Identification of the FANCI protein, a monoubiquitinated FANCD2 paralog required for DNA repair. Cell. 2007;129(2):289–301 https://doi.org/10.1016/j.cell.2007.03.009.

16. Montes de Oca R, Andreassen PR, Margossian SP, Gregory RC, Taniguchi T, Wang X, et al. Regulated interaction of the Fanconi anemia protein, FANCD2, with chromatin. Blood. 2005;105(3):1003–9 https://doi.org/10.1182/blood-2003-11-3997.

17. Longerich S, Kwon Y, Tsai MS, Hlaing AS, Kupfer GM, Sung P. Regulation of FANCD2 and FANCI monoubiquitination by their interaction and by DNA. Nucleic Acids Res. 2014;42(9):5657–70 https://doi.org/10.1093/nar/gku198.

18. Liang C-C, Li Z, Lopez-Martinez D, Nicholson WV, Vénien-Bryan C, Cohn MA. The FANCD2–FANCI complex is recruited to DNA interstrand crosslinks before monoubiquitination of FANCD2. Nature Communications. 2016;7:12124 https://doi.org/10.1038/ncomms12124 https://www.nature.com/articles/ncomms12124#supplementary-information.

19. Joo W, Xu G, Persky NS, Smogorzewska A, Rudge DG, Buzovetsky O, et al. Structure of the FANCI-FANCD2 Complex: Insights into the Fanconi Anemia DNA Repair Pathway. Science. 2011;333(6040):312 https://doi.org/10.1126/science.1205805.

20. Longerich S, Kwon Y, Tsai M-S, Hlaing AS, Kupfer GM, Sung P. Regulation of FANCD2 and FANCI monoubiquitination by their interaction and by DNA. Nucleic acids research. 2014;42(9):5657–70 https://doi.org/10.1093/nar/gku198.

21. Longerich S, San Filippo J, Liu D, Sung P. FANCI Binds Branched DNA and Is Monoubiquitinated by UBE2T-FANCL. Journal of Biological Chemistry. 2009;284(35):23182–6

22. Niraj J, Caron M-C, Drapeau K, Bérubé S, Guitton-Sert L, Coulombe Y, et al. The identification of FANCD2 DNA binding domains reveals nuclear localization sequences. Nucleic acids research. 2017;45(14):8341–57 https://doi.org/10.1093/nar/gkx543.

23. van Twest S, Murphy VJ, Hodson C, Tan W, Swuec P, O’Rourke JJ, et al. Mechanism of Ubiquitination and Deubiquitination in the Fanconi Anemia Pathway. Mol Cell. 2017;65(2):247–59 https://doi.org/10.1016/j.molcel.2016.11.005.

24. Tan W, Murphy VJ, Charron A, van Twest S, Sharp M, Bythell-Douglas R, et al. Preparation and purification of mono-ubiquitinated proteins using Avi-tagged ubiquitin. in preparation. 2019

25. Kratz K, Schöpf B, Kaden S, Sendoel A, Eberhard R, Lademann C, et al. Deficiency of FANCD2-Associated Nuclease KIAA1018/FAN1 Sensitizes Cells to Interstrand Crosslinking Agents. Cell. 2010;142(1):77–88 https://doi.org/10.1016/j.cell.2010.06.022.

26. Wojtaszek JL, Wang S, Kim H, Wu Q, D’Andrea AD, Zhou P. Ubiquitin recognition by FAAP20 expands the complex interface beyond the canonical UBZ domain. Nucleic Acids Res. 2014;42(22):13997–4005 https://doi.org/10.1093/nar/gku1153.

27. Castillo A, Paul A, Sun B, Huang Ting H, Wang Y, Yazinski Stephanie A, et al. The BRCA1-Interacting Protein Abraxas Is Required for Genomic Stability and Tumor Suppression. Cell Reports. 2014;8(3):807–17 https://doi.org/10.1016/j.celrep.2014.06.050.

28. Densham RM, Garvin AJ, Stone HR, Strachan J, Baldock RA, Daza-Martin M, et al. Human BRCA1-BARD1 ubiquitin ligase activity counteracts chromatin barriers to DNA resection. Nat Struct Mol Biol. 2016;23(7):647–55 https://doi.org/10.1038/nsmb.3236 http://www.nature.com/nsmb/journal/v23/n7/abs/nsmb.3236.html#supplementary-information.

29. Raghunandan M, Chaudhury I, Kelich SL, Hanenberg H, Sobeck A. FANCD2, FANCJ and BRCA2 cooperate to promote replication fork recovery independently of the Fanconi Anemia core complex. Cell cycle (Georgetown, Tex). 2015;14(3):342–53 https://doi.org/10.4161/15384101.2014.987614.

30. Jacquemont C, Taniguchi T. Proteasome Function Is Required for DNA Damage Response and Fanconi Anemia Pathway Activation. Cancer Res. 2007;67(15):7395

31. Lossaint G, Larroque M, Ribeyre C, Bec N, Larroque C, Décaillet C, et al. FANCD2 Binds MCM Proteins and Controls Replisome Function upon Activation of S Phase Checkpoint Signaling. Mol Cell. 2013;51(5):678–90 https://doi.org/10.1016/j.molcel.2013.07.023.

32. Wang B, Matsuoka S, Ballif BA, Zhang D, Smogorzewska A, Gygi SP, et al. Abraxas and RAP80 Form a BRCA1 Protein Complex Required for the DNA Damage Response. Science. 2007;316(5828):1194

33. Yang K, Moldovan G-L, D’Andrea AD. RAD18-dependent Recruitment of SNM1A to DNA Repair Complexes by a Ubiquitin-binding Zinc Finger. J Biol Chem. 2010;285(25):19085–91

34. van Twest S, Murphy VJ, Hodson C, Tan W, Swuec P, O’Rourke JJ, et al. Mechanism of Ubiquitination and Deubiquitination in the Fanconi Anemia Pathway. Mol Cell. 2017;65(2):247–59 https://doi.org/10.1016/j.molcel.2016.11.005.

35. Sareen A, Chaudhury I, Adams N, Sobeck A. Fanconi anemia proteins FANCD2 and FANCI exhibit different DNA damage responses during S-phase. Nucleic acids research. 2012;40(17):8425–39 https://doi.org/10.1093/nar/gks638.

36. Thompson EL, Yeo JE, Lee E-A, Kan Y, Raghunandan M, Wiek C, et al. FANCI and FANCD2 have common as well as independent functions during the cellular replication stress response. Nucleic acids research. 2017;45(20):11837–57 https://doi.org/10.1093/nar/gkx847.

37. Husnjak K, Dikic I. Ubiquitin-binding proteins: decoders of ubiquitin-mediated cellular functions. Annu Rev Biochem. 2012;81:291–322 https://doi.org/10.1146/annurev-biochem-051810-094654.

38. Swuec P, Renault L, Borg A, Shah F, Murphy VJ, van Twest S, et al. The FA Core Complex Contains a Homo-dimeric Catalytic Module for the Symmetric Mono-ubiquitination of FANCI-FANCD2. Cell reports. 2017;18(3):611–23 https://doi.org/10.1016/j.celrep.2016.11.013.

39. Liang CC, Li Z, Lopez-Martinez D, Nicholson WV, Venien-Bryan C, Cohn MA. The FANCD2-FANCI complex is recruited to DNA interstrand crosslinks before monoubiquitination of FANCD2. Nat Commun. 2016;7:12124 https://doi.org/10.1038/ncomms12124.

40. Lopez-Martinez D, Kupculak M, Yang D, Yoshikawa Y, Liang CC, Wu R, et al. Phosphorylation of FANCD2 Inhibits the FANCD2/FANCI Complex and Suppresses the Fanconi Anemia Pathway in the Absence of DNA Damage. Cell reports. 2019;27(10):2990–3005.e5 https://doi.org/10.1016/j.celrep.2019.05.003.

41. Wang AT, Smogorzewska A. SnapShot: Fanconi Anemia and Associated Proteins. Cell. 2015;160(1-2):354–e1 https://doi.org/10.1016/j.cell.2014.12.031.

42. Gregory RC, Taniguchi T, D’Andrea AD. Regulation of the Fanconi anemia pathway by monoubiquitination. Semin Cancer Biol. 2003;13(1):77–82

43. Tian Y, Shen X, Wang R, Klages-Mundt NL, Lynn EJ, Martin SK, et al. Constitutive role of the Fanconi anemia D2 gene in the replication stress response. J Biol Chem. 2017;292(49):20184–95 https://doi.org/10.1074/jbc.M117.814780.

44. Madireddy A, Kosiyatrakul ST, Boisvert RA, Herrera-Moyano E, Garcia-Rubio ML, Gerhardt J, et al. FANCD2 Facilitates Replication through Common Fragile Sites. Mol Cell. 2016;64(2):388–404 https://doi.org/10.1016/j.molcel.2016.09.017.

45. MacKay C, Declais AC, Lundin C, Agostinho A, Deans AJ, MacArtney TJ, et al. Identification of KIAA1018/FAN1, a DNA repair nuclease recruited to DNA damage by monoubiquitinated FANCD2. Cell. 2010;142(1):65–76 https://doi.org/10.1016/j.cell.2010.06.021.

46. Rego MA, Kolling FWt, Vuono EA, Mauro M, Howlett NG. Regulation of the Fanconi anemia pathway by a CUE ubiquitin-binding domain in the FANCD2 protein. Blood. 2012;120(10):2109–17 https://doi.org/10.1182/blood-2012-02-410472.

47. Zhang Y. Transcriptional regulation by histone ubiquitination and deubiquitination. Genes Dev. 2003;17(22):2733–40 https://doi.org/10.1101/gad.1156403.

48. Greer SF, Zika E, Conti B, Zhu X-S, Ting JPY. Enhancement of CIITA transcriptional function by ubiquitin. Nature Immunology. 2003;4(11):1074–82 https://doi.org/10.1038/ni985.

49. Alcón P, Shakeel S, Patel KJ, Passmore LA. FANCD2-FANCI is a clamp stabilized on DNA by monoubiquitination during DNA repair. bioRxiv. 2019:858225 https://doi.org/10.1101/858225.

50. Wang R, Wang S, Dhar A, Peralta C, Pavletich NP. DNA clamp function of the monoubiquitinated Fanconi Anemia FANCI-FANCD2 complex. bioRxiv. 2019:854133 https://doi.org/10.1101/854133.

51. Deans AJ, West SC. FANCM connects the genome instability disorders Bloom’s Syndrome and Fanconi Anemia. Mol Cell. 2009;36(6):943–53 https://doi.org/10.1016/j.molcel.2009.12.006.

52. Iacovoni JS, Caron P, Lassadi I, Nicolas E, Massip L, Trouche D, et al. High-resolution profiling of gammaH2AX around DNA double strand breaks in the mammalian genome. Embo j. 2010;29(8):1446–57 https://doi.org/10.1038/emboj.2010.38.

53. Sato K, Ishiai M, Toda K, Furukoshi S, Osakabe A, Tachiwana H, et al. Histone chaperone activity of Fanconi anemia proteins, FANCD2 and FANCI, is required for DNA crosslink repair. EMBO J. 2012;31(17):3524–36 https://doi.org/10.1038/emboj.2012.197.

54. Higgs MR, Sato K, Reynolds JJ, Begum S, Bayley R, Goula A, et al. Histone Methylation by SETD1A Protects Nascent DNA through the Nucleosome Chaperone Activity of FANCD2. Mol Cell. 2018;71(1):25–41.e6 https://doi.org/10.1016/j.molcel.2018.05.018.

55. Lim CJ, Lee SY, Teramoto J, Ishihama A, Yan J. The nucleoid-associated protein Dan organizes chromosomal DNA through rigid nucleoprotein filament formation in E. coli during anoxia. Nucleic Acids Res. 2013;41(2):746–53 https://doi.org/10.1093/nar/gks1126.

56. Hui MP, Galkin VE, Yu X, Stasiak AZ, Stasiak A, Waldor MK, et al. ParA2, a Vibrio cholerae chromosome partitioning protein, forms left-handed helical filaments on DNA. Proc Natl Acad Sci U S A. 2010;107(10):4590–5 https://doi.org/10.1073/pnas.0913060107.

57. Misra HS, Maurya GK, Chaudhary R, Misra CS. Interdependence of bacterial cell division and genome segregation and its potential in drug development. Microbiological research. 2018;208:12–24 https://doi.org/10.1016/j.micres.2017.12.013.

58. de Laat WL, Appeldoorn E, Sugasawa K, Weterings E, Jaspers NG, Hoeijmakers JH. DNA-binding polarity of human replication protein A positions nucleases in nucleotide excision repair. Genes Dev. 1998;12(16):2598–609 https://doi.org/10.1101/gad.12.16.2598.

59. Chen H, Lisby M, Symington LS. RPA coordinates DNA end resection and prevents formation of DNA hairpins. Mol Cell. 2013;50(4):589–600 https://doi.org/10.1016/j.molcel.2013.04.032.

60. Nguyen HD, Yadav T, Giri S, Saez B, Graubert TA, Zou L. Functions of Replication Protein A as a Sensor of R Loops and a Regulator of RNaseH1. Mol Cell. 2017;65(5):832–47.e4 https://doi.org/10.1016/j.molcel.2017.01.029.

61. Neelsen KJ, Lopes M. Replication fork reversal in eukaryotes: from dead end to dynamic response. Nat Rev Mol Cell Biol. 2015;16(4):207–20 https://doi.org/10.1038/nrm3935.

62. Thangavel S, Berti M, Levikova M, Pinto C, Gomathinayagam S, Vujanovic M, et al. DNA2 drives processing and restart of reversed replication forks in human cells. J Cell Biol. 2015;208(5):545–62 https://doi.org/10.1083/jcb.201406100.

63. Sidorova JM, Kehrli K, Mao F, Monnat R, Jr. Distinct functions of human RECQ helicases WRN and BLM in replication fork recovery and progression after hydroxyurea-induced stalling. DNA Repair (Amst). 2013;12(2):128–39 https://doi.org/10.1016/j.dnarep.2012.11.005.

64. Richardson CD, Kazane KR, Feng SJ, Bray NL, Schaefer AJ, Floor S, et al. CRISPR-Cas9 Genome Editing In Human Cells Works Via The Fanconi Anemia Pathway. bioRxiv. 2017:doi: https://doi.org/10.1101/136028

65. Sato K, Shimomuki M, Katsuki Y, Takahashi D, Kobayashi W, Ishiai M, et al. FANCI-FANCD2 stabilizes the RAD51-DNA complex by binding RAD51 and protects the 5’-DNA end. Nucleic Acids Res. 2016;44(22):10758–71 https://doi.org/10.1093/nar/gkw876.

66. Dubois EL, Guitton-Sert L, Beliveau M, Parmar K, Chagraoui J, Vignard J, et al. A Fanci knockout mouse model reveals common and distinct functions for FANCI and FANCD2. Nucleic Acids Res. 2019;47(14):7532–47 https://doi.org/10.1093/nar/gkz514.

67. Thompson EL, Yeo JE, Lee EA, Kan Y, Raghunandan M, Wiek C, et al. FANCI and FANCD2 have common as well as independent functions during the cellular replication stress response. Nucleic Acids Res. 2017;45(20):11837–57 https://doi.org/10.1093/nar/gkx847.

68. Sareen A, Chaudhury I, Adams N, Sobeck A. Fanconi anemia proteins FANCD2 and FANCI exhibit different DNA damage responses during S-phase. Nucleic Acids Res. 2012;40(17):8425–39 https://doi.org/10.1093/nar/gks638.

69. Castella M, Jacquemont C, Thompson EL, Yeo JE, Cheung RS, Huang JW, et al. FANCI Regulates Recruitment of the FA Core Complex at Sites of DNA Damage Independently of FANCD2. PLoS Genet. 2015;11(10):e1005563 https://doi.org/10.1371/journal.pgen.1005563.

70. Yang K, Moldovan GL, Vinciguerra P, Murai J, Takeda S, D’Andrea AD. Regulation of the Fanconi anemia pathway by a SUMO-like delivery network. Genes Dev. 2011;25(17):1847–58 https://doi.org/10.1101/gad.17020911.

71. Butturini A, Gale RP, Verlander PC, Adler-Brecher B, Gillio AP, Auerbach AD. Hematologic abnormalities in Fanconi anemia: an International Fanconi Anemia Registry study. Blood. 1994;84(5):1650–5

72. Soulier J, Leblanc T, Larghero J, Dastot H, Shimamura A, Guardiola P, et al. Detection of somatic mosaicism and classification of Fanconi anemia patients by analysis of the FA/BRCA pathway. Blood. 2005;105(3):1329–36 https://doi.org/10.1182/blood-2004-05-1852.

73. de la Rosa-Trevín JM, Otón J, Marabini R, Zaldívar A, Vargas J, Carazo JM, et al. Xmipp 3.0: An improved software suite for image processing in electron microscopy. Journal of Structural Biology. 2013;184(2):321–8 https://doi.org/https://doi.org/10.1016/j.jsb.2013.09.015.

74. Mindell JA, Grigorieff N. Accurate determination of local defocus and specimen tilt in electron microscopy. Journal of Structural Biology. 2003;142(3):334–47 https://doi.org/https://doi.org/10.1016/S1047-8477(03)00069-8.

75. Grant T, Rohou A, Grigorieff N. cisTEM, user-friendly software for single-particle image processing. eLife. 2018;7:e35383 https://doi.org/10.7554/eLife.35383.

