## Supplementary tables for "Monoubiquitination by the Fanconi Anemia core complex locks FANCI:FANCD2 on DNA in filamentous arrays"

**Supplementary materials**

Supplementary Table 1. DNA oligonucleotides used in this study.

The following oligo nucleotides were ordered from Integrated DNA Technologies (Singapore):

| **Oligo No.** | **Sequence** |
| --- | --- |
| XOM1 | 5’-ACGCTGCCGAATTCTACCAGTGCCTT  GCTAGGACATCTTTGCCCACCTGCAGG  TTCACCC-3' |
| XOM2 | 5’-GGGTGAACCTGCAGGTGGGCAAAGA  TGTCCATCTGTTGTAATCGTCAAGCTTTA  TGCCGT-3' |
| XOM3 | 5’-ACGGCATAAAGCTTGACGATTACAACA  GATCATGGAGCTGTCTAGAGGATCCGAC  TATCG-3’ |
| XOM4 | 5’-CGATAGTCGGATCCTCTAGACAGCTCC  ATGTAGCAAGGCACTGGTAGAATTCGGCA  GCGT-3’ |
| XOM2.1/2 | 5’-GGGTGAACCTGCAGGTGGGCAAAGATG  TCC-3’ |
| XOM3.1/2 | 5’-CATGGAGCTGTCTAGAGGATCCGACTAT  CG-3’ |
| XOM1.comp | 5'-GGGTGAACCTGCAGGTGGGCAAAGATGT  CCTAGCAAGGCACTGGTAGAATTCGGCAGCGT-3' |
| oligo1-150 | 5'-TAAATAAGATAAGGATAATACAAAATAAGTA  AATGAATAAACAGAGAAAATAAAGTAAAGGAT  ATAAAAAATGAACATAAAGAATAAGTAAATGAA  TAAAACATAATAGGAATAAATATAGGAAATGAA  ATAAAAGAGACATAAATAAGA-3' |
| oligo2-150 | 5'-TCTTATTTATGTCTCTTTTATTTCATTTCCTATAT  TTATTCCTATTATGTTTTATTCATTTACTTATTCTTT  ATGTTCATTTTTTATATCCTTTACTTTATTTTCTCTG  TTTATTCATTTACTTATTTTGTATTATCCTTATCTTA  TTTA-3' |

Supplementary Table 2. Combination of oligonucleotides annealed to generate DNA substrates used in this study.

| **Substrate** | **labelled oligo** | **mix with cold oligo** |
| --- | --- | --- |
| ssDNA | 12 µM XOM1 | - |
| dsDNA | 12 µM XOM1 | 36 µM XOM1.com |
| splayed arm | 12 µM XOM1 | 36 µM XOM4 |
| 3' flap | 12 µM XOM1 | 36 µM XOM3.1/2 |
|  |  | 36 µM XOM4 |
| 5' flap | 12 µM XOM1 | 36 µM XOM2.1/2 |
|  |  | 36 µM XOM4 |
| replication fork | 12 µM XOM1 | 36 µM XOM2.1/2 |
|  |  | 36 µM XOM3.1/2 |
|  |  | 36 µM XOM4 |
| 150 bp dsDNA |  | 12 uM oligo1-150 |
|  |  | 12 uM oligo2-150 |
